# Modeling traumatic brain injury combined with hemorrhagic shock in rats: Neurological assessment and PET imaging with ^18^F-fluorodeoxyglucaric acid

**DOI:** 10.1101/2021.06.01.446662

**Authors:** Hibah O. Awwad, Andria Hedrick, Alex Mdzinarishvili, Hailey Houson, Kelly M. Standifer, Vibhudutta Awasthi

**Author notes:** **Corresponding Authors:** Hibah O. Awwad, Ph.D., Department of Pharmaceutical Sciences, College of Pharmacy, University of Oklahoma Health Sciences Center, 1110 N. Stonewall Ave. CPB Ste 315, Oklahoma City, OK 73117, Vibhudutta Awasthi, Ph.D., Department of Pharmaceutical Sciences, College of Pharmacy, University of Oklahoma Health Sciences Center, 1110 N. Stonewall Ave. CPB Ste 309, Oklahoma City, OK 73117.

## Abstract

Traumatic brain injury (TBI) is a major cause of death and disability worldwide. Hemorrhagic shock (HS) aggravates tissue injury and complicates TBI recovery. We studied the combined insult of mild TBI and HS and investigated the impact of varying loss of blood volume on neurologic deficit and brain lesion volume. A novel positron emission tomography (PET) technique was employed to monitor tissue injury. Male Sprague Dawley rats received mTBI by controlled cortical impact (CCI) followed by withdrawal of 0%, 30-40%, 45%, or 50% of blood (mTBI, mTBI+HS_≤40%_, mTBI+HS_45%_, and mTBI+HS_50%_, respectively). Neurological deficit (mNSS= 5.6, 7.6, and 12.3) and mortality (2/12, 2/6, and 7/12) were worse in mTBI+HS_≤40%_, mTBI+HS_45%_, and mTBI+HS_50%_, respectively than in mTBI alone rats (no death; mNSS=3.3). Histologic lesion size increased 3.5-fold in mTBI+HS_50%_ compared to mTBI alone and the infarct-avid PET agent ^18^F-fluorodeoxyglucaric acid (FGA) proportionately detected tissue necrosis in mTBI+HS_50%_ rats. Based on these results, we conclude that HS aggravates mTBI-induced neurological deficits, tissue injury and mortality. PET/^18^F-FGA as an imaging marker can detect the extent of injury in a non-invasive manner.

## INTRODUCTION

Traumatic brain injury (TBI) is the leading cause of death and disability in young adults below 40 years of age in the United States; similar epidemiological statistics exists worldwide^1, 2^. A TBI is a blow or jolt to the head by an external force resulting in primary injury damage due to mechanical disruption, axonal injury and necrosis of brain tissue, followed by a delayed secondary injury, which collectively manifests as inflammation, neurodegeneration and neurobehavioral deficits^3–5^. The immediate primary injury from a TBI results in focal damage due to mechanical shear stress and presents with tissue necrosis, cortical contusions, vascular injury, and intracranial hemorrhage accompanied by cerebral vasospasms. Brain ischemia, edema and neuroinflammation characterize the subsequent, delayed secondary injury^4, 5^. Recovery from a TBI depends on the severity of the damage, timing of intervention, and the presence of co-morbid conditions or associated pathophysiological events^3, 6^. For instance, a TBI incident can be accompanied by significant blood loss (e.g. severe hemorrhage from a peripheral wound), resulting in an injury accentuated by hemorrhagic shock (HS) ^7–9^.

Clinical HS is classified into four classes, depending on the percent of blood volume lost: Classes I, II, III, and IV correspond to <15%, 15-30%, 30-40% and >40% blood loss, respectively^10, 11^. Both HS and TBI independently contribute to the development of systemic inflammatory response syndrome ^7–9, 12^. When HS is associated with TBI, the usual compensatory shift of blood flow from splanchnic organs to the brain is ineffective in restoration of cerebral function because of hypovolemia and hypotension^13^. Data from the traumatic Coma Databank and preclinical literature confirm that hypotension and hypoxemia are independently associated with significant increases in morbidity and mortality from severe head injury^6, 7, 11^. The larger the amount of blood loss, the more aggravating is the impact of HS on TBI-associated impairments in cerebral metabolism, redox status, ion homeostasis, and decoupling of cerebral blood flow from metabolic demand^7–9^. Mortality in TBI cases also increases exponentially with severity of blood loss^14^. Mild TBI (mTBI) is the most common grade of TBI, but its association with HS has not been intensively investigated. To address this gap of great medical significance, we studied the effect of HS on mTBI in a rat model.

This article describes important hemodynamic, neurological and survival characteristics of a polytrauma rat model of mTBI±HS in which a controlled cortical impact (CCI)-induced mTBI was followed by graded withdrawal of blood to induce various levels of HS. The conventional CCI is the most commonly used experimental TBI model that is very suitable for understanding the pathology of mTBI complicated by concomitant HS. The controlled impact with the electromagnetic piston to the exposed dura results in translatable neurological, behavioral and biochemical consequences similar to those observed in TBI patients^15 16^. We also introduce a positron emission tomography (PET) technique using a novel infarct imaging marker ^18^F-fluorodeoxyglucaric acid (FGA) to monitor the severity of the mTBI+HS injury by non-invasive imaging.

## MATERIALS AND METHODS

### Animals

Male (n=51) Sprague-Dawley rats (200-250 g) were purchased from Envigo Laboratories (Houston, TX) and were acclimated for 1 week upon receipt. Housed in pairs, rats received *ad libitum* access to food and water and a 12 h dark/light cycle. The University of Oklahoma Health Sciences Center (OUHSC) Institutional Animal Care and Use Committee and the USAMRDC Animal Care and Use Review Office (ACURO) approved all animal studies. Experiments were conducted and are reported in compliance with ARRIVE guidelines^17^. Sample size was not determined *a priori* as this study was the first to evaluate the effect of mTBI±HS on rat survival rates, mNSS, and use of PET to assess severity of injury following polytrauma. Rats were randomized after cannulation surgery using the ‘=RAND()’ function in Microsoft Excel. Block randomization was performed to reduce variability in the average weights on the day of polytrauma. Rats were anesthetized for all surgical procedures with 3% isoflurane in medical grade air delivered at 800 mL/min. Rats were continuously monitored and vital signs were recorded during anesthesia and surgery. Twenty-four hours post-mTBI±HS, rats were euthanized by injecting 150 μL Somnasol® (Henry Schein, NY). The same person performed all animal surgeries; all personnel assessing outcomes were blind to the treatment groups. Exclusion criterion included more than 10% blood loss during TBI or cannulation surgeries, blocked arterial catheter, impaired limb movement, and baseline neurological score following cannulation > 6. The sequence of procedures, from femoral artery cannulation to euthanasia, is shown in Fig. 1.

**Figure 1:**
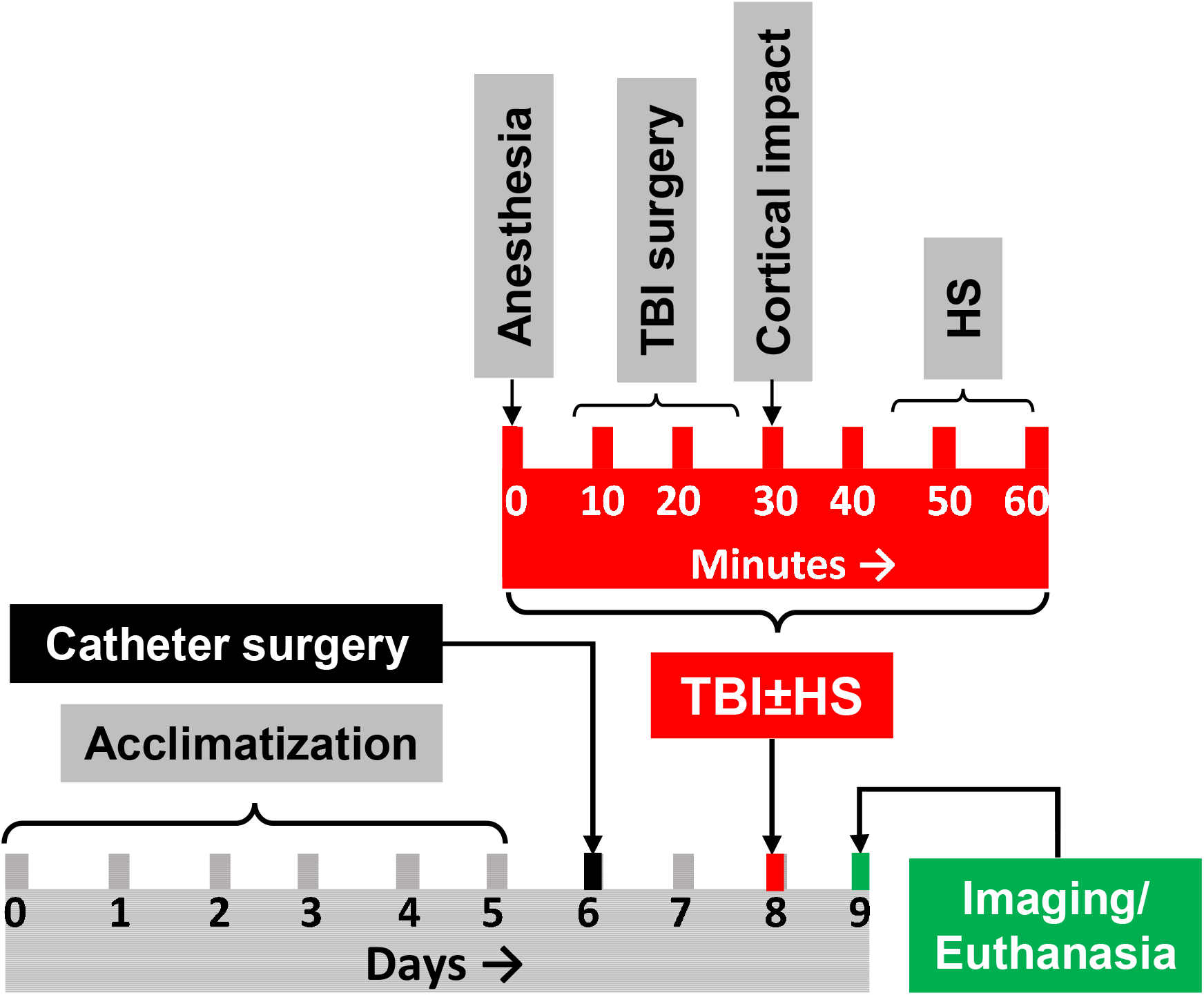
Sequence of procedures implemented on rats recruited in mTBI±HS study. After 5 days of acclimatization, femoral artery catheter was implanted. The rats were given 2 days to recover before mTBI±HS. The mTBI±HS step lasted approximately 1 h. The rats were allowed to recover without any respiratory or thermal support and monitored for 24 h. Afterwards, the rats were euthanized. Selected rats were also subjected to imaging study prior to euthanasia.

### Femoral artery cannulation

Indwelling catheters were implanted in the left femoral artery of each rat under isoflurane-medical air anesthesia and aseptic conditions, as described previously^18^. The catheter was tunneled under the skin to the nape of neck and secured with a suture. Catheter patency was maintained with a heparin block (1 kU/mL). Rats received a single subcutaneous dose of buprenorphine SR (1 mg/kg) for post-surgical analgesia.

### Mild TBI using controlled cortical impact (CCI)

Mild TBI was induced by a standard protocol for CCI using the electromagnetic Impact One device (Leica Biosystems, IL)^19, 20^. Briefly, anesthetized rats stabilized in a stereotaxic frame received a midline scalp incision. Craniotomy (~6 mm diameter) exposed the intact dura mater over the point of impact (3 mm left lateral and 1.8 mm posterior to Bregma). Using digital dual manipulators for precision, a 2 mm diameter tip TBI Impact One actuator angled at 20° from vertical position impacted the brain at a velocity of 3 m/s and dwelling time of 100 ms to a 4 mm depth below the dura surface. After mTBI induction, the craniotomy site was sealed with bone wax, the skin was glued with tissue adhesive, triple antibiotic ointment was applied to the wound. Time to righting reflex and rat movement was assessed post-anesthesia. Rats in the sham group differed from mTBI rats only in that they did not receive an impact to the brain.

### Hemorrhagic shock

HS was performed approximately 15-30 min after rats recovered from mTBI surgery anesthesia by withdrawing blood via the arterial catheter as described previously^21–23^. Following the constant volume model, varying volumes of blood (0%, 35-40%, 44-46%, and 48-50% of total blood) were withdrawn at a rate of 1.0 ml/min to generate four trauma groups: mTBI alone, mTBI+HS_≤40%_, mTBI+HS_45%_, and mTBI+HS_50%_, respectively. The total blood volume was estimated as 7% of body weight. Following HS, the rats were allowed to recover from anesthesia without any additional support. None of the surviving animals showed signs of distress that required early euthanasia or exclusion from the study.

### Mean arterial pressure (MAP) and heart rate (HR)

MAP, HR, and rectal temperature were recorded at three different time points: baseline (pre-HS or 0 h), 6 h post-HS, and 24 h post-HS. MAP and HR were measured directly from the femoral artery catheter by a BP-102 transducer coupled with ETH-225 Bio-amplifier, and 118 analog-to-digital converter (iWorx, Dover, NH). The data were recorded while rats were under isoflurane anesthesia; MAP and HR were calculated from the recordings using Labscribe 3.0 software (iWorx).

### Neurological function assessment

Rat neurological status was determined using a modified neurological severity score (mNSS)^24^. Baseline assessments were performed 48 h post-cannulation and immediately before mTBI or sham craniotomy surgeries on day 0 (D0). Postinjury scores were assessed one day (D1) after mTBI±HS injury. Neurological deficits are designated as mild, moderate, or severe based on mNSS scores in the range of 1-6, 7-12, or 13-18, respectively.^24^ Rats with intact neurological function received scores ranging from 0 to <1. Non-surviving rats following polytrauma received the highest D1 mNSS of 18. One rat with mNSS score > 6 prior to mTBI procedure were excluded from the study due to complications related to the femoral artery cannulation.

### PET Imaging with ^18^F-labeled fluorodeoxyglucaric acid (^18^F-FGA)

For PET imaging, ^18^F-FGA was synthesized from commercially available ^18^F-fluorodeoxyglucose (^18^F-FDG). The methods of synthesis and quality control are described in previous reports^25, 26^. Anesthetized rats received approximately 1 mCi of ^18^F-FGA intravenously via the tail vein. The tracer was allowed to distribute for 2 h before imaging. List-mode data were acquired for 20 min, followed by a computed tomogram (CT) using a Flex X-O/X-PET/CT scanner (Gamma Medica-Ideas, Northridge, CA). In some cases, ^18^F-FGA uptake was also monitored by *ex vivo* imaging of extracted whole brains from sham, mTBI, and mTBI+HS_50%_ groups. After image reconstruction, the PET and CT images were fused and ^18^F-FGA uptake in ipsilateral and contralateral hemispheres was calculated by drawing 3-dimensional regions of interest (ROI) by Amira software. The uptake values were normalized to the injected radioactive dose (ID) after correcting for background and decay as needed. This PET/CT imaging was accomplished at the Research Imaging Facility in the OUHSC-College of Pharmacy (Oklahoma City, OK).

To validate imaging, the ipsilateral and contralateral hemispheres were also counted for radioactivity at necropsy. An automated gamma counter (Perkin Elmer, MA) was used for tissue counting and the tissue associated counts were expressed as counts per min per gram of tissue (CPM/g).

### Enzyme-linked immunoassay (ELISA) for plasma biomarkers

Blood samples were collected in heparinized tubes and plasma was separated by centrifugation at 5,000 rpm for 5 min at 4°C. The samples collected 24 h after mTBI+HS were stored at −80°C until assayed for interleukin-6 (IL-6), tumor necrosis factor-α (TNF-α), nerve growth factor (NGF), and brain-derived neurotrophic factor (BDNF). The manufacturer’s recommendations for the respective rat ELISA kits were followed (Bosterbio, Pleasanton, CA). Duplicate samples were normalized to protein concentration. We used 5x, 1x, 1x, and 200x dilutions for IL-6, TNF-α, NGF, and BDNF, respectively.

### Histopathology and lesion volume measurement

Triphenyltetrazolium chloride (TTC)-staining utilized 2 mm-thick sections from freshly extracted brain 24 h after mTBI±HS. Briefly, the coronal sections were stained at 37 °C for 20 min and digital pictures of the stained sections were captured. Lesion area and whole brain area were traced from each TTC-stained slice image using NIH Image J (version 1.52a; NIH). Measurements were set to scale in mm using a ruler captured in the same field of view and multiplied by the thickness of each slice (2 mm) to calculate lesion volume as percentage of the whole brain volume^27^.

### Statistical analysis

Data are presented as mean ± standard deviation. In this study, n refers to the number of animals or the number of samples collected (one from each animal). Data analysis employed an appropriate statistical test as indicated in the text using GraphPad Prism v.9. Differences were considered significant if p < 0.05. Raw data were used from all studies except where indicated; outliers were removed by the Grubbs’ method with an alpha of 0.05.

## RESULTS

The polytrauma model of CCI-induced mTBI with various degrees of HS in male Sprague-Dawley rats was employed to interrogate the contribution of HS to mTBI. Mild TBI conditions were kept constant, but the amount of blood withdrawn was varied to induce HS with ≤40%, 45%, or 50% blood loss. Physiological and neurological changes resulting from multiple levels of mTBI±HS polytrauma and in sham TBI rats were assessed over a 24 h period following HS.

### Mean arterial pressure and heart rate

Continuous blood pressure monitoring during the blood withdrawal phase characterized the anticipated hypovolemia. Baseline MAP did not differ between groups and mTBI alone had no significant effect on MAP values compared to baseline (Fig. 2A). As expected, MAP dropped immediately following mTBI+HS_≤40%_, mTBI+HS_45%_, and mTBI+HS_50%_ compared to mTBI alone (*p<0.0001) and mTBI+HS_40%_ groups (#p<0.01). MAP recovery in mTBI+HS_≤40%_, a class III HS, was statistically indistinguishable from mTBI alone. Mild TBI with class IV HS groups recovered more slowly; both back to baseline at 24 h. However, mTBI+HS_45%_ group showed MAP values significantly lower than mTBI alone (Fig. 2A) at 6 hr (*p<0.05). Analysis with a mixed effects 2-way ANOVA model (REML) found a significant interaction between time and polytrauma [F(_2, 30_)=36; p<0.0001]. Significant effects of time (p<0.0001) and polytrauma (p=0.0008) were noted. Most rats died between HS and the 6 hr time point, especially in the mTBI+HS_50%_ group. Since each time point reflects surviving rats at that time, MAP values appear to return to baseline by 6 hr for that group. For HR, modest effects of time [F(2, 47)=5.651; p=0.0063] and polytrauma [F(3, 47)=3.78; p=0.0165] were found with the 2 way-ANOVA analysis, but no significant interaction between time and polytrauma or post-hoc differences were noted (Fig. 2B). Rectal temperature ranged from 35-38 °C in various groups and no differences was observed between groups.

**Figure 2:**
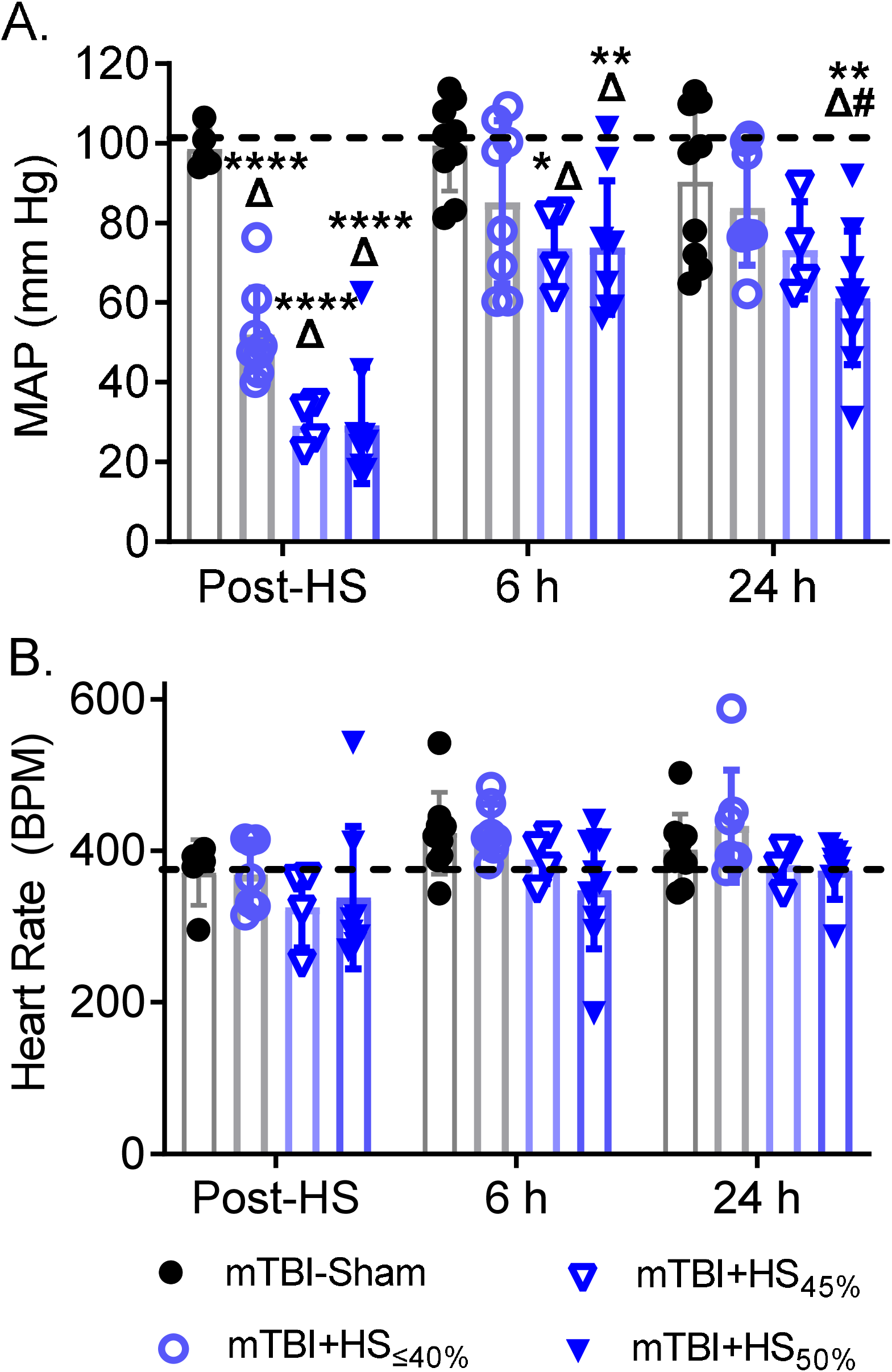
Hemodynamic profile of rats with mTBI and varying degree of HS. (A) Mean arterial pressure (MAP) and (B) heart rate data were recorded at baseline, immediately after HS (post-HS), 6 h post-HS and 24 h post-HS. The baseline MAP (101.4±1.3 mm Hg) and HR (391±27 BPM) are shown as horizontal dashed lines (*p<0.0001 vs. baseline, ^Δ^p<0.05 vs. mTBI alone, #p<0.05 vs. mTBI+HS_≤40%_).

### Neurological function assessment

Severity of mTBI was determined using the mNSS test 24 h after mTBI±HS and compared to mNSS on D0 (Table 1). Femoral artery cannulation, which was performed 48 h before performing mTBI or sham craniotomy surgery, produced a mild deficit in pre-mTBI D0 mNSS (mNSS=1.1 ± 0.6, n=51), but it did not differ significantly from normal healthy rats (mNSS=1.0 ± 1.8, n=9). Rats with indwelling catheters exhibited restricted movement over the first two days; all rats were included in the final analysis except for two rats with pre-mTBI D0 mNSS scores > 6. Rats in the mTBI alone exhibited D1 mNSS values ranging between 1 and 6, consistent with a mild neurological deficit. The mNSS values increased with volume of blood loss, indicating increased severity of neurological deficit (Table 1). Accordingly, mTBI+HS_≤40%_, mTBI+HS_45%_ and mTBI+HS_50%_ groups displayed moderate-severe neurological deficits, with mNSS values ranging between 1 and 18. The most severe neurological deficits were noted in mTBI+HS_50%_, with mNSS values of 12.0 ± 7.4. Two-way ANOVA revealed significant interaction between the time factor and polytrauma condition [F (_4, 45_)=6.954; p=0.0002], a significant effect of time [F(_1, 45_)=38.13; p<0.0001], and a significant effect of trauma [F (_4, 45_)=4.572; p=0.0035]. Post-hoc comparisons with Sidak’s multiple comparison test indicated significant increase in neurological deficits only in mTBI+HS_50%_ compared to their D0 control values (*p<0.0001). We also calculated ΔmNSS as the change in mNSS for each group between days 0 and 1. One-way ANOVA for ΔmNSS values also mirrored the severity of injury with a significant effect of polytrauma [F4, 45)=6.954; p= 0.0002]; post-hoc analysis revealed that mTBI+HS_50%_ differed from Sham control (*p<0.001), from mTBI alone (^ΔΔΔ^p<0.001) and from TBI+≤HS_40%_ (##p<0.01) groups.

**Table 1:** Neurological function assessment of mTBI+HS rats. Rats were scored based on mNSS assessment before mTBI/Sham surgery (Day 0) and 24 h after mTBI+HS (Day 1). Data are presented as mean ± S.D. (*p<0.05 vs control and ^Δ^mTBI alone).

| Assessment | Modified neurological severity score (mNSS) |  |  |  |  |
| --- | --- | --- | --- | --- | --- |
|  | Control | mTBI alone | mTBI+HS<sub>≤40%</sub> | mTBI+HS<sub>45%</sub> | mTBI+HS<sub>50%</sub> |
| Day 0 (D0) | 0.8 ± 1.6 | 1.2 ± 1.6 | 1.6 ± 1.7 | 1.7 ± 1.4 | 0.4 ± 2.6 |
| Day 1 (D1) | 1.6 ± 1.9 | 3.3 ± 2.7 | 5.6 ± 6.3 | 7.58 ± 8.2 | **12.3 ± 7.0\*\*\*** |
| #rats/group | n=9 | n=11 | n=12 | n=6 | n=12 |
| ΔmNSS<br>(D1 minus D0) | 0.8 ± 1.6 | 2.2 ± 3.1 | 4.0 ± 5.6 | 5.9 ± 7.6 | **12.0±7.4\*\*\*,ΔΔ,##** |

### Survival (Table 2)

**Table 2:** Survival rate (24 h) and mNSS of surviving rats. measured 24 h post-mTBI±HS (Day 1). Data presented as mean ± S.D. Treatment groups significantly differed for survival (p=0.0002) by Chi-Square test for trend. The mNSS score in surviving rats did not differ between groups by one-way ANOVA with Tukey’s post-hoc test.

| Parameter | Control | mTBI alone | mTBI+HS<sub>≤40%</sub> | mTBI+HS<sub>45%</sub> | mTBI+HS<sub>50%</sub> |
| --- | --- | --- | --- | --- | --- |
| # Died /Total* | 0/9 | 0/11 | 2/12 | 2/6 | 7/12 |
| % Survival | 100% | 100 % | 83.3% | 66.7% | 58.3% |
| mNSS | 1.6 ± 2.0 | 3.3 ± 2.7 | 3.1 ± 2.8 | 2.4 ± 1.6 | 4.4 ± 0.8 |

All rats in the sham surgery (control) group and mTBI alone groups survived. The mortality occurred only with the polytrauma (mTBI+HS), confirming that HS is an aggravating factor in mTBI. Survival rates declined with increasing severity of HS (p=0.0002) by Chi-square test for trend). It is important to note that even with the polytrauma condition of mTBI+HS_50%_, which caused the highest mortality rate, the surviving rats displayed mild neurological deficits on day post-injury that did not differ between groups.

### Histology

Tissue injury was evaluated by determining lesion volumes from direct morphometry of formalin-fixed, unstained tissue sections (Fig. 3A). TTC staining (Fig. 3B) of mTBI-induced necrotic lesions (~1-2 mm in size) confirmed injury to the left sensorimotor cortex and hippocampus. The necrotic lesion volume (unstained area) at the injury site increased relative to the amount of blood withdrawn. The most extensive necrosis was in the mTBI+HS_50%_ group. As shown in Fig. 3C, groups mTBI+HS_40%_ and mTBI+HS_45%_ produced lesion volumes that were similar to the lesion volume produced by mTBI alone (approximately 3-4% of the brain volume). However, lesion volume in the most severe polytrauma group, mTBI+HS_50%_, increased 3.5-fold and constituted 10.7% of the brain volume (one-way ANOVA, p<0.05).

**Figure 3:**
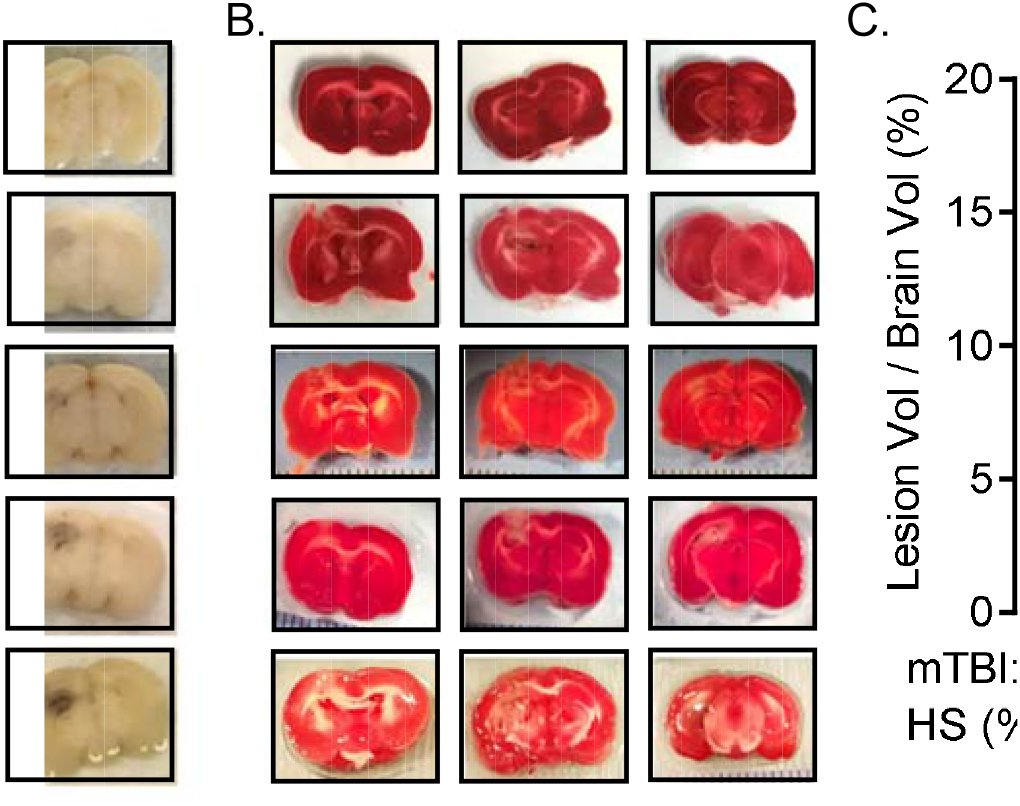
HS results in an increase in the mTBI lesion size. (A) Representative pictures of formalin fixed brain sections. (B) Lesion volumes were calculated by tracing the respective areas in each formalin-fixed slice and expressed as a % of whole brain volume. (C) TTC-stained sections of brain for each treatment group. Data are presented as mean ±S.D. (*p < 0.05 vs every other treatment using one-way ANOVA; n=3-4/group).

### PET imaging of polytrauma-induced lesions with ^18^F-FGA

Since significant differences in lesion size were only found in mTBI+HS_50%_, live animal PET imaging studies were performed to compare FGA accumulation in mTBI and mTBI+HS_50%_ groups (Fig. 4). CT images showed physical damage to the skull due to craniotomy (arrow; Fig. 4A). PET images showed ipsilateral accumulation of ^18^F-FGA in mTBI+HS_50%_ brain (green), but there was no brain uptake in mTBI alone rat (Fig. 4B). Image analysis by ROI drawing on ipsilateral and contralateral hemispheres indicated significant difference in quantified values of ^18^F-FGA uptake ratios between the two treatment groups (Fig. 4C).

**Figure 4:**
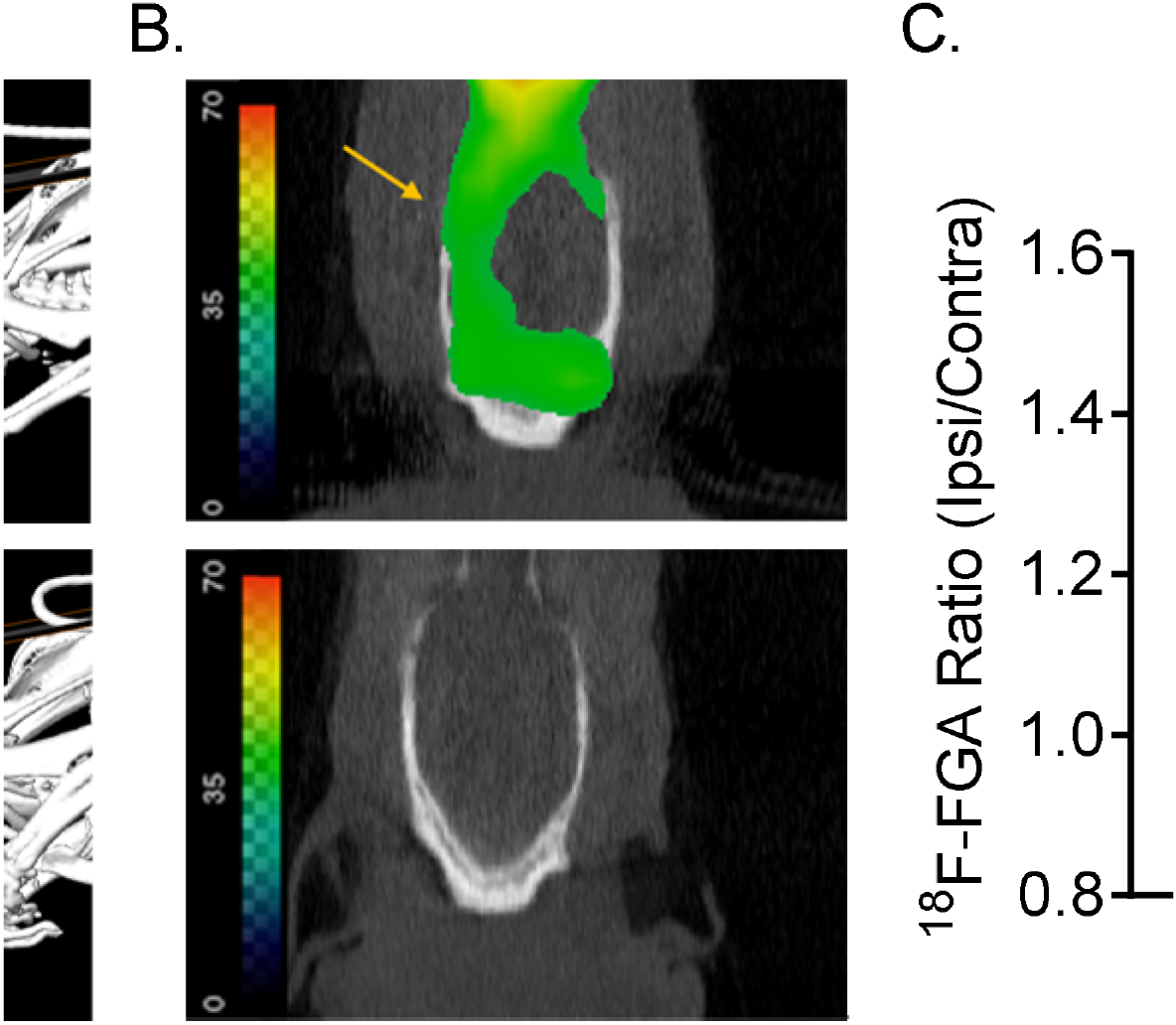
Visualization of mTBI+HS injury by PET/^18^F-FGA. (A) CT images of cranial field of view. Arrow points to the location of the cortical impact. The rat’s head was supported by a stand to obtain proper orientation with respect to the horizontal plane. Also shown is the plane of the digital slice used to derive coronal PET images in XY-plane. (B) Longitudinal section of PET tomograms along the digital slice on CT images. Increased ^18^F-FGA uptake is shown in the ipsilateral (left) hemisphere in mTBI+HS_50%_ rat. (C) Quantification of ^18^F-FGA uptake in ipsilateral and contralateral hemispheres in mTBI alone and mTBI+HS_50%_ groups, presented as a ratio (n=5 each). The digital data was obtained by drawing ROIs as described in methods and compared using an unpaired student’s t-test.

^18^F-FGA accumulation was also quantified based on *ex vivo* images of extracted brains (Fig. 5). The brain images showed clear differences in ^18^F-FGA uptake among sham, mTBI, and mTBI+HS_50%_ rats (Fig. 5A). Figure 5B shows TTC-stained brain coronal slices of an mTBI+HS_50%_ rat recruited in imaged study. In the mTBI+HS_50%_ group, the contralateral and ipsilateral brain tissue showed significant difference in radioactive counts (Fig. 5C). ^18^F-FGA accumulated 5-fold more in the ipsilateral tissue as compared to the accumulation in the contralateral non-injured brain tissue (p<0.0001). Together, these imaging results corroborated the histological and neurological findings that HS increases the severity of brain injury caused by localized mTBI, and indicated that PET imaging could be employed as a sensitive and non-invasive procedure to assess TBI-induced brain injury.

**Figure 5:**
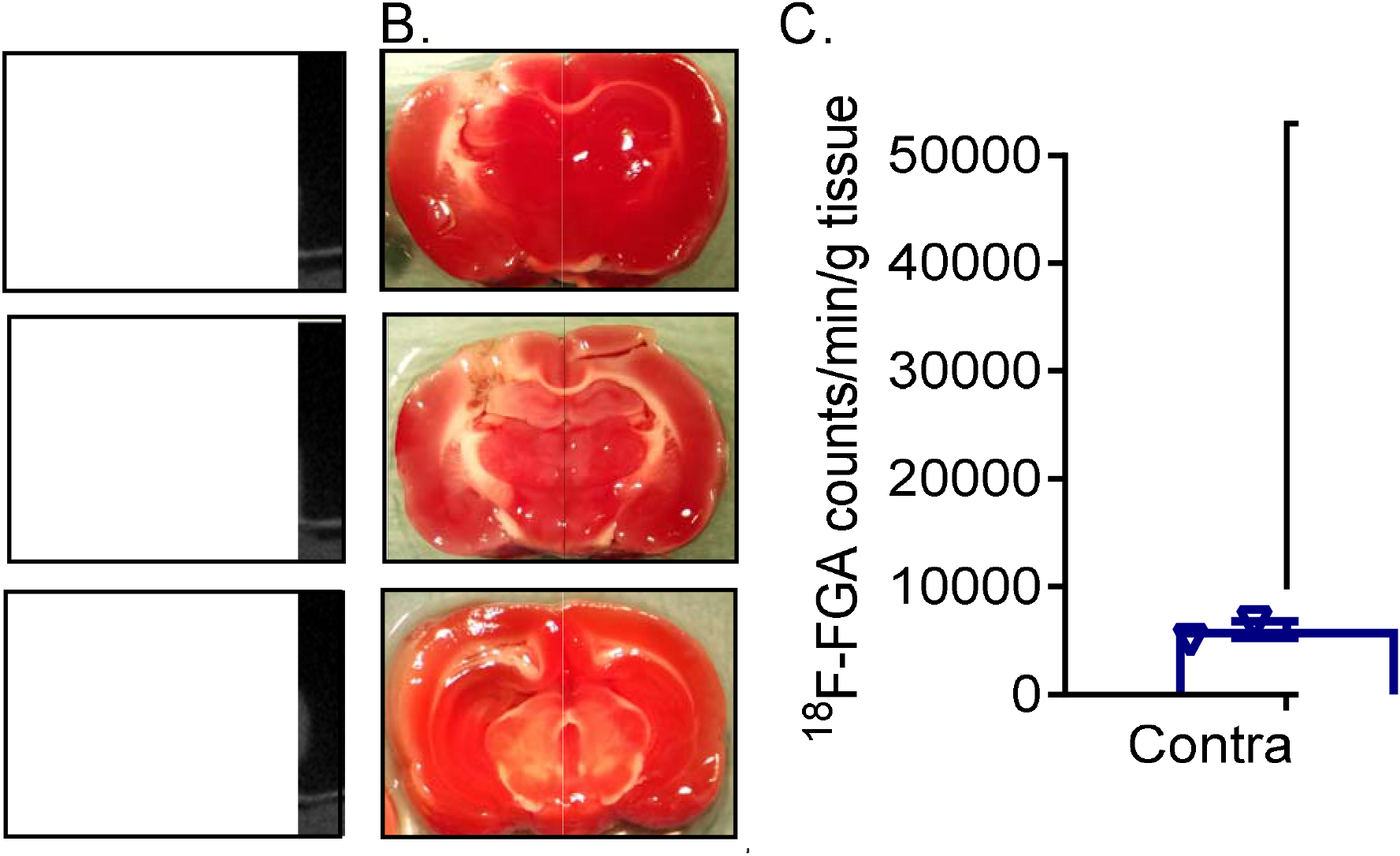
^18^F-FGA accumulates in ipsilateral brain tissue of rats with mTBI±HS injury. (A) PET images of excised whole brain. (B) TTC-stained coronal brain slices from mTBI+HS_50%_ showing tissue damage. (C) Radioactive counts associated with contralateral and ipsilateral tissue of mTBI+HS_50%_ rats (*p < 0.0001). The data are expressed as counts per min per gram of tissue and compared using an unpaired student’s t-test.

### Neurotrophins and inflammatory cytokines in blood plasma

Levels of BDNF, NGF, IL-6, and TNF-α were determined in blood plasma collected at baseline and 24 h after polytrauma. There was a significant effect of polytrauma on plasma BDNF levels collected 24 h post-trauma [F(3,13)=11.2; p=0.007], with over 2-fold increases in BDNF in the mTBI+HS_≤40%_ group (Fig. 6); the increase in mTBI+HS_≤40%_ was significant compared to mTBI alone (p<0.01), mTBI+HS_45%_ (p<0.05),, and mTBI+HS_50%_ (p<0.001) as determined by one-way ANOVA analysis with Tukey’s multiple comparison. No significant differences were noted in 24 h IL-6 levels compared to baseline or between trauma groups (data not shown); TNF-α and NGF concentrations were below the detection limit of the ELISA in the plasma samples tested.

**Figure 6:**
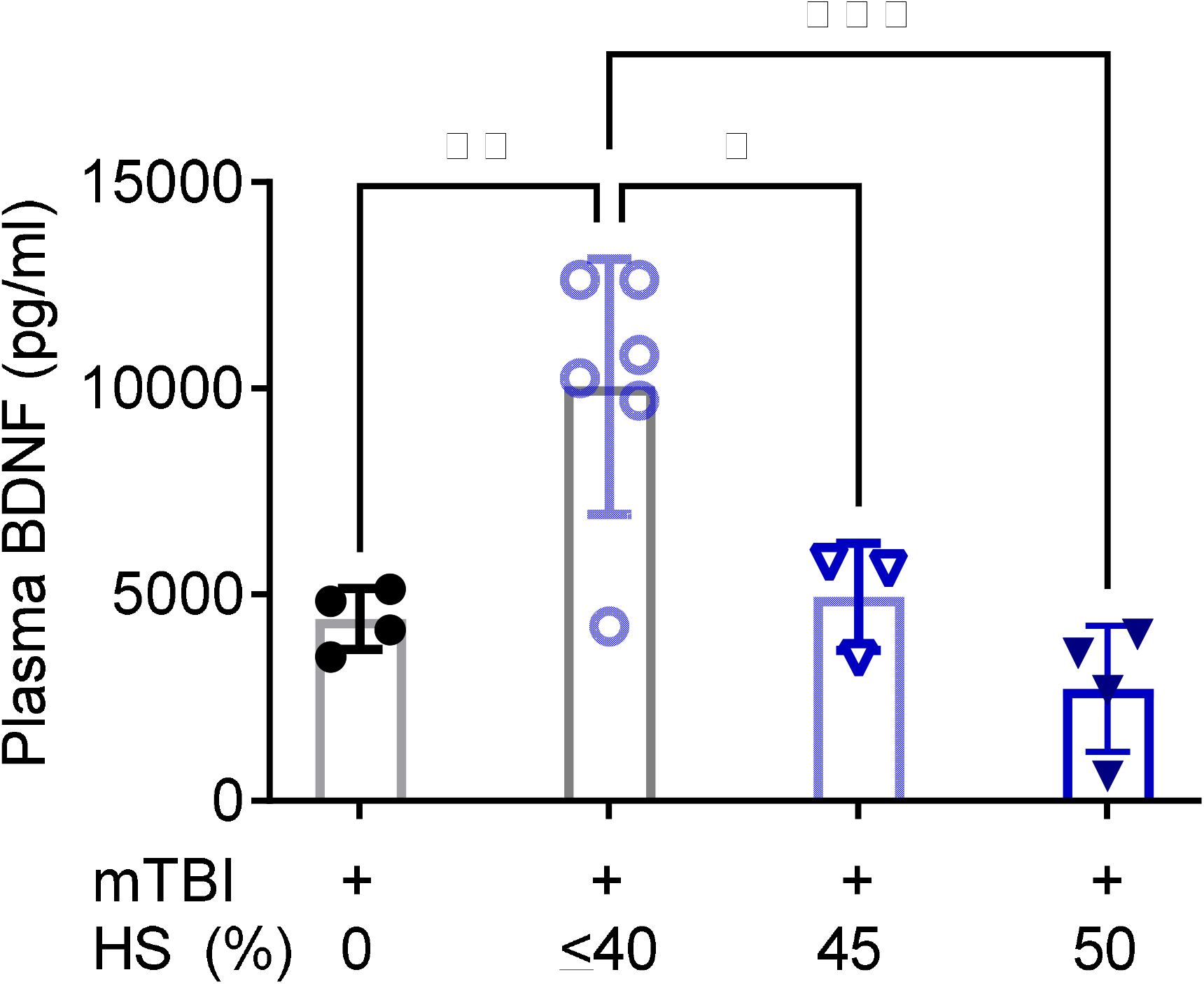
Plasma BDNF levels in 24 h samples. Data are presented as scatter plot overlay of bar graphs that show mean ± S.D.

## DISCUSSION

This study expands the current understanding of experimental models of TBI accompanied by HS that have been reported previously^8, 9, 28–30^. We assessed the effect of varying degrees of HS on mTBI-associated neurological outcome and tissue injury in the brain. We also investigated a non-invasive PET technique to monitor extent of tissue injury in mTBI+HS model. We focused on creating mild TBI as it constitutes 90% of the TBI cases identified in modern conflicts (Operation Iraqi Freedom/Operation Endurance Freedom^31–33^; similar statistics are found among civilians^34, 35^. Combined mTBI+HS injury is frequently observed in automobile accidents and military conflicts. Since the majority of deaths from such polytrauma injury occur within the first 24 h, this study characterized the consequences during the first 24 h of mTBI+HS.

Hypotension secondary to the blood loss is an expected outcome in hypovolemic shock^8, 13^, but it also influences local brain injury caused by trauma in a complex and paradoxical manner. There are conflicting reports on the effect of HS alone on brain damage. Yamauchi, et al. found hippocampal damage after HS in rats^36^, but Carrillo, et al. reported that prolonged HS and resuscitation in a rat model failed to cause functional or histological changes in the brain^37^. In contrast, combined TBI+HS pathology is consistently more severe than TBI alone. Chesnut et al. reported that hypovolemia and hypoxia increased TBI-induced morbidity by 150%^6^. HS influences brain injury by causing ischemia, bioenergetic imbalance^21^, and promotion of excitatory signaling in the brain^38, 39^. The aggravated TBI+HS tissue injury clinically manifests as increased mortality or severe neurological deficits^6, 7, 40, 41^. In our rat model, the most severe neurological deficits were experienced by rats exposed to class IV HS (>45% blood loss). Twenty four h mortality was also high in mTBI+HS_45%_ and mTBI+HS_50%_ groups. However, mTBI+HS_45-50%_ rats that survived beyond 6 h of polytrauma displayed intact physiological reflexes, complete hemodynamic recovery, and only mild neurological deficits. Moreover, rats belonging to mTBI+HS_≤40%_ and mTBI+HS_50%_ groups showed more rapid recovery, perhaps due to reduction in intracranial pressure (ICP) secondary to hypovolemia in the rats that ultimately survived. Elevated ICP is an aggravating factor in the injured brain as white matter and brainstem compression following elevated ICP may produce death from respiratory depression^42, 43^. We hypothesize that at a certain threshold of hypovolemia, benefits of reduced ICP outweigh the risk associated with combined injury. Reduced blood volume relieves potential damage caused by brain edema and inflammation that otherwise would cause the brain tissue to swell. This explanation is supported by a study from Hariri, et al. who reported that crystalloid resuscitation elevated central venous pressure by reducing brain compliance, and thus worsened the outcome in a swine model^44^. Therefore, it is important to manage fluid resuscitation in patients with critical brain injury^45, 46^.

Another outcome from our study was the PET technique using ^18^F-FGA to assess brain necrotic lesions. Importantly, the image signal due to ^18^F-FGA accumulation was proportional to the extent of histologically detected brain damage. The volume of the mTBI-induced lesion detected by PET/^18^F-FGA increased significantly in mTBI+HS_50%_ rats. Since imaging did not show substantial change in brain ^18^F-FGA accumulation at 44-46% blood loss, it appears that the threshold for impact of HS on mTBI lies between 47% and 50% blood loss. Clinical application of this imaging technique can accurately evaluate the size of a necrotic lesion, which will be helpful in assessing both prognosis and recovery of a patient following TBI±HS. Although molecular targets of FGA in necrotic lesions are not clear yet, it is believed to bind to the nuclear proteins, such as histones, which are exposed in cells undergoing necrosis^25^. To date, PET imaging in TBI is limited to the use of ^18^F-FDG to image glucose metabolism^47, 48^. Unlike ^18^F-FDG, ^18^F-FGA does not accumulate in normal healthy brain tissue, which would otherwise create difficulties in interpretation of images. Direct visualization of the necrotic core is mechanistically different from other radiologic techniques of CT and magnetic resonance imaging (MRI) as they depend on delayed structural or morphologic changes^49^. In contrast, physiological changes, including initiation of necrotic cell death due to oxidative stress, metabolic dysfunction, inflammation and excitotoxicity occur relatively early.

Clinical management of TBI+HS mostly depends on vital signs, hemodynamic monitoring, and plasma biomarkers. Among plasma-based biomarkers studied in this work, only BDNF levels significantly varied by polytrauma. TNF-α and NGF were below the detection limit of the assay, while IL-6 levels did not differ among groups (not shown). Higher plasma BDNF levels were found at 24 h in mTBI+HS_≤40%_ rats, a group that exhibited neither significant neurological deficits, nor reduced survival. However, the increase in plasma BDNF levels was not apparent with higher levels of blood loss. The absence of inflammatory cytokines in plasma of mTBI+HS groups was unexpected, especially when both TBI and HS independently contribute to the pathogenesis of systemic inflammatory response syndrome^50–52^. Inflammatory mediators also contribute to early, delayed, and systemic effects of TBI^53^. Literature review provided no explanation for this unusual finding, but one report suggested that the reduced blood volume could have induced an anti-inflammatory response in the presence of TBI^54^. It is also possible that cytokine upregulation due to combined injury occurred prior to the 6 h time point and subsided before 24 h, thus missing the optimal time of sample collection.

## CONCLUSIONS

Overall, we conclude that mild TBI with 50% blood loss is the optimal polytrauma model for combined injury caused by HS and mTBI in rats. Below 50% blood loss, the aggravating impact of HS on brain injury is not consistent. At the same time, it is important to note that within a short dynamic range of 45-50% blood loss, 24 h outcomes such as MAP, brain lesion size, and neurological deficits rapidly worsen. We also report the effective use of PET/^18^F-FGA to image TBI in proportion to the extent of brain injury. PET imaging with FGA has a potential to impact diagnostic assessment of polytrauma TBI+HS injury in a clinical setting, including for the evaluation of sports-related concussion and accidental head injury.

## ACKNOWLEDGEMENTS

Authors acknowledge Cindy Simpson Durand and Megan Lerner for their technical contributions to this project. This work was supported by the Department of the Army-Prolonged Field Care Research Award #W81XWH-16-DMRDP-CCCRP-PFCRA to Dr. Vibhudutta Awasthi.

## AUTHOR CONTRIBUTION STATEMENT

HOA, KMS, and VA designed the study and analyzed the data.

HOA, VA, AH, and ML performed the experiments and/or collected the data.

VA, KMS and HOA wrote the manuscript. All authors have read the manuscript, provided their feedback, and approved the submission of the final version

## DISCLOSURE/CONFLICT OF INTEREST

*The authors declare that the research was conducted in the absence of any commercial or financial relationships that could be construed as a potential conflict of interest*.

## Notes

### Competing Interest Statement

The authors have declared no competing interest.

## REFERENCES

1. Thurman DJ, Alverson C, Dunn KA, et al. Traumatic brain injury in the United States: A public health perspective. J Head Trauma Rehabil 1999; 14: 602–615. 2000/02/15. DOI: 10.1097/00001199-199912000-00009.

2. Taylor CA, Bell JM, Breiding MJ, et al. Traumatic Brain Injury-Related Emergency Department Visits, Hospitalizations, and Deaths - United States, 2007 and 2013. MMWR Surveill Summ 2017; 66: 1–16. 2017/03/17. DOI: 10.15585/mmwr.ss6609a1.

3. Moppett IK. Traumatic brain injury: assessment, resuscitation and early management. Br J Anaesth 2007; 99: 18–31. 2007/06/05. DOI: 10.1093/bja/aem128.

4. Gaetz M. The neurophysiology of brain injury. Clin Neurophysiol 2004; 115: 4–18. 2004/01/07. DOI: 10.1016/s1388-2457(03)00258-x.

5. McAllister TW. Neurobiological consequences of traumatic brain injury. Dialogues Clin Neurosci 2011; 13: 287–300. 2011/10/29.

6. Chesnut RM, Marshall LF, Klauber MR, et al. The role of secondary brain injury in determining outcome from severe head injury. The Journal of trauma 1993; 34: 216–222. Research Support, U.S. Gov’t, P.H.S. 1993/02/01.

7. Galvagno SM, Jr., Fox EE, Appana SN, et al. Outcomes after concomitant traumatic brain injury and hemorrhagic shock: A secondary analysis from the Pragmatic, Randomized Optimal Platelets and Plasma Ratios trial. J Trauma Acute Care Surg 2017; 83: 668–674. 2017/09/21. DOI: 10.1097/TA.0000000000001584.

8. Leung LY, Deng-Bryant Y, Shear D, et al. Experimental Models Combining TBI, Hemorrhagic Shock, and Hypoxemia. Methods Mol Biol 2016; 1462: 445–458. 2016/09/09. DOI: 10.1007/978-1-4939-3816-2_25.

9. Vogt N, Herden C, Roeb E, et al. Cerebral Alterations Following Experimental Multiple Trauma and Hemorrhagic Shock. Shock 2018; 49: 164–173. 2017/07/07. DOI: 10.1097/SHK.0000000000000943.

10. Mutschler M, Paffrath T, Wolfl C, et al. The ATLS((R)) classification of hypovolaemic shock: a well established teaching tool on the edge? Injury 2014; 45 Suppl 3: S35–38. 2014/10/07. DOI: 10.1016/j.injury.2014.08.015.

11. Cannon JW. Hemorrhagic Shock. N Engl J Med 2018; 378: 370–379. 2018/01/25. DOI: 10.1056/NEJMra1705649.

12. Chakraborty RK and Burns B. Systemic Inflammatory Response Syndrome. StatPearls.Treasure Island (FL), 2020.

13. DeWitt DS and Prough DS. Blast-induced brain injury and posttraumatic hypotension and hypoxemia. Journal of neurotrauma 2009; 26: 877–887. Review 2008/05/02. DOI: 10.1089/neu.2007.0439.

14. Doppenberg EM, Choi SC and Bullock R. Clinical trials in traumatic brain injury: lessons for the future. Journal of neurosurgical anesthesiology 2004; 16: 87–94. Review 2003/12/17.

15. Bondi CO, Semple BD, Noble-Haeusslein LJ, et al. Found in translation: Understanding the biology and behavior of experimental traumatic brain injury. Neurosci Biobehav Rev 2015; 58: 123–146. 2014/12/17. DOI: 10.1016/j.neubiorev.2014.12.004.

16. Dhillon HS, Donaldson D, Dempsey RJ, et al. Regional levels of free fatty acids and Evans blue extravasation after experimental brain injury. Journal of neurotrauma 1994; 11: 405–415. 1994/08/01. DOI: 10.1089/neu.1994.11.405.

17. Percie du Sert N, Hurst V, Ahluwalia A, et al. The ARRIVE guidelines 2.0: updated guidelines for reporting animal research. J Physiol 2020; 598: 3793–3801. 2020/07/16. DOI: 10.1113/JP280389.

18. Awasthi V, Yee SH, Jerabek P, et al. Cerebral oxygen delivery by liposome-encapsulated hemoglobin: a positron-emission tomographic evaluation in a rat model of hemorrhagic shock. J Appl Physiol (1985) 2007; 103: 28–38. 2007/07/07. DOI: 10.1152/japplphysiol.00136.2006.

19. Osier ND, Korpon JR and Dixon CE. Controlled Cortical Impact Model. In: Kobeissy FH (ed) Brain Neurotrauma: Molecular, Neuropsychological, and Rehabilitation Aspects. Boca Raton (FL), 2015.

20. Brody DL, Mac Donald C, Kessens CC, et al. Electromagnetic controlled cortical impact device for precise, graded experimental traumatic brain injury. Journal of neurotrauma 2007; 24: 657–673. 2007/04/19. DOI: 10.1089/neu.2006.0011.

21. Rao G, Xie J, Hedrick A, et al. Hemorrhagic shock-induced cerebral bioenergetic imbalance is corrected by pharmacologic treatment with EF24 in a rat model. Neuropharmacology 2015; 99: 318–327. 2015/08/02. DOI: 10.1016/j.neuropharm.2015.07.033.

22. Yadav VR, Hussain A, Sahoo K, et al. Remediation of hemorrhagic shock-induced intestinal barrier dysfunction by treatment with diphenyldihaloketones EF24 and CLEFMA. J Pharmacol Exp Ther 2014; 351: 413–422. 2014/09/11. DOI: 10.1124/jpet.114.217331.

23. Yadav VR, Sahoo K, Roberts PR, et al. Pharmacologic suppression of inflammation by a diphenyldifluoroketone, EF24, in a rat model of fixed-volume hemorrhage improves survival. J Pharmacol Exp Ther 2013; 347: 346–356. 2013/09/03. DOI: 10.1124/jpet.113.208009.

24. Chen J, Sanberg PR, Li Y, et al. Intravenous administration of human umbilical cord blood reduces behavioral deficits after stroke in rats. Stroke 2001; 32: 2682–2688. Research Support, Non-U.S. Gov’t 2001/11/03.

25. Houson HA, Nkepang GN, Hedrick AF, et al. Imaging of isoproterenol-induced myocardial injury with (18)F labeled fluoroglucaric acid in a rat model. Nuclear medicine and biology 2018; 59: 9–15. 2018/02/08. DOI: 10.1016/j.nucmedbio.2017.12.006.

26. Houson H, Mdzinarishvili A, Gali H, et al. PET Detection of Cerebral Necrosis Using an Infarct-Avid Agent 2-Deoxy-2-[(18)F]Fluoro-D-Glucaric Acid (FGA) in a Mouse Model of the Brain Stroke. Mol Imaging Biol 2020 2020/06/20. DOI: 10.1007/s11307-020-01513-9.

27. Elliott MB, Jallo JJ and Tuma RF. An investigation of cerebral edema and injury volume assessments for controlled cortical impact injury. J Neurosci Methods 2008; 168: 320–324. 2007/12/14. DOI: 10.1016/j.jneumeth.2007.10.019.

28. Navarro JC, Pillai S, Cherian L, et al. Histopathological and behavioral effects of immediate and delayed hemorrhagic shock after mild traumatic brain injury in rats. Journal of neurotrauma 2012; 29: 322–334. 2011/11/15. DOI: 10.1089/neu.2011.1979.

29. Salci K, Enblad P, Goiny M, et al. Metabolic effects of a late hypotensive insult combined with reduced intracranial compliance following traumatic brain injury in the rat. Ups J Med Sci 2010; 115: 221–231. 2010/10/28. DOI: 10.3109/03009734.2010.503906.

30. Robertson CS, Cherian L, Shah M, et al. Neuroprotection with an erythropoietin mimetic peptide (pHBSP) in a model of mild traumatic brain injury complicated by hemorrhagic shock. Journal of neurotrauma 2012; 29: 1156–1166. 2011/05/07. DOI: 10.1089/neu.2011.1827.

31. Shin SS, Bales JW, Edward Dixon C, et al. Structural imaging of mild traumatic brain injury may not be enough: overview of functional and metabolic imaging of mild traumatic brain injury. Brain imaging and behavior 2017; 11: 591–610. Review 2017/02/15. DOI: 10.1007/s11682-017-9684-0.

32. Hoge CW, McGurk D, Thomas JL, et al. Mild traumatic brain injury in U.S. Soldiers returning from Iraq. N Engl J Med 2008; 358: 453–463. 2008/02/01. DOI: 10.1056/NEJMoa072972.

33. Ling GS and Ecklund JM. Traumatic brain injury in modern war. Curr Opin Anaesthesiol 2011; 24: 124–130. 2011/02/09. DOI: 10.1097/ACO.0b013e32834458da.

34. Gardner RC and Yaffe K. Epidemiology of mild traumatic brain injury and neurodegenerative disease. Mol Cell Neurosci 2015; 66: 75–80. 2015/03/10. DOI: 10.1016/j.mcn.2015.03.001.

35. Cassidy JD, Carroll LJ, Peloso PM, et al. Incidence, risk factors and prevention of mild traumatic brain injury: results of the WHO Collaborating Centre Task Force on Mild Traumatic Brain Injury. J Rehabil Med 2004: 28–60. 2004/04/16. DOI: 10.1080/16501960410023732.

36. Yamauchi Y, Kato H and Kogure K. Hippocampal damage following repeated brief hypotensive episodes in the rat. J Cereb Blood Flow Metab 1991; 11: 974–978. 1991/11/01. DOI: 10.1038/jcbfm.1991.163.

37. Carrillo P, Takasu A, Safar P, et al. Prolonged severe hemorrhagic shock and resuscitation in rats does not cause subtle brain damage. The Journal of trauma 1998; 45: 239-248; discussion 248-239. 1998/08/26. DOI: 10.1097/00005373-199808000-00007.

38. Guerriero RM, Giza CC and Rotenberg A. Glutamate and GABA imbalance following traumatic brain injury. Curr Neurol Neurosci Rep 2015; 15: 27. 2015/03/23. DOI: 10.1007/s11910-015-0545-1.

39. Yamauchi Y, Kato H and Kogure K. Brain damage in a new hemorrhagic shock model in the rat using long-term recovery. J Cereb Blood Flow Metab 1990; 10: 207–212. 1990/03/01. DOI: 10.1038/jcbfm.1990.36.

40. Petrosoniak A and Hicks C. Resuscitation Resequenced: A Rational Approach to Patients with Trauma in Shock. Emerg Med Clin North Am 2018; 36: 41–60. 2017/11/15. DOI: 10.1016/j.emc.2017.08.005.

41. Dennis AM, Haselkorn ML, Vagni VA, et al. Hemorrhagic shock after experimental traumatic brain injury in mice: effect on neuronal death. Journal of neurotrauma 2009; 26: 889–899. 2008/09/11. DOI: 10.1089/neu.2008.0512.

42. Friess SH, Lapidus JB and Brody DL. Decompressive craniectomy reduces white matter injury after controlled cortical impact in mice. Journal of neurotrauma 2015; 32: 791–800. 2015/01/06. DOI: 10.1089/neu.2014.3564.

43. Shen L, Wang Z, Su Z, et al. Effects of Intracranial Pressure Monitoring on Mortality in Patients with Severe Traumatic Brain Injury: A Meta-Analysis. PLoS One 2016; 11: e0168901. 2016/12/29. DOI: 10.1371/journal.pone.0168901.

44. Hariri RJ, Firlick AD, Shepard SR, et al. Traumatic brain injury, hemorrhagic shock, and fluid resuscitation: effects on intracranial pressure and brain compliance. J Neurosurg 1993; 79: 421–427. 1993/09/01. DOI: 10.3171/jns.1993.79.3.0421.

45. van der Jagt M. Fluid management of the neurological patient: a concise review. Crit Care 2016; 20: 126. 2016/06/01. DOI: 10.1186/s13054-016-1309-2.

46. Nevin DG and Brohi K. Permissive hypotension for active haemorrhage in trauma. Anaesthesia 2017; 72: 1443–1448. 2017/09/25. DOI: 10.1111/anae.14034.

47. Byrnes KR, Wilson CM, Brabazon F, et al. FDG-PET imaging in mild traumatic brain injury: a critical review. Front Neuroenergetics 2014; 5: 13. 2014/01/11. DOI: 10.3389/fnene.2013.00013.

48. Awwad HO, Gonzalez LP, Tompkins P, et al. Blast Overpressure Waves Induce Transient Anxiety and Regional Changes in Cerebral Glucose Metabolism and Delayed Hyperarousal in Rats. Front Neurol 2015; 6: 132. 2015/07/03. DOI: 10.3389/fneur.2015.00132.

49. Mutch CA, Talbott JF and Gean A. Imaging Evaluation of Acute Traumatic Brain Injury. Neurosurg Clin N Am 2016; 27: 409–439. 2016/09/18. DOI: 10.1016/j.nec.2016.05.011.

50. Weaver LC, Bao F, Dekaban GA, et al. CD11d integrin blockade reduces the systemic inflammatory response syndrome after traumatic brain injury in rats. Exp Neurol 2015; 271: 409–422. 2015/07/15. DOI: 10.1016/j.expneurol.2015.07.003.

51. Namas R, Ghuma A, Hermus L, et al. The acute inflammatory response in trauma / hemorrhage and traumatic brain injury: current state and emerging prospects. Libyan J Med 2009; 4: 97–103. 2009/01/01. DOI: 10.4176/090325.

52. Mi Q, Constantine G, Ziraldo C, et al. A dynamic view of trauma/hemorrhage-induced inflammation in mice: principal drivers and networks. PLoS One 2011; 6: e19424. 2011/05/17. DOI: 10.1371/journal.pone.0019424.

53. Lu J, Goh SJ, Tng PY, et al. Systemic inflammatory response following acute traumatic brain injury. Front Biosci (Landmark Ed) 2009; 14: 3795–3813. 2009/03/11. DOI: 10.2741/3489.

54. Shein SL, Shellington DK, Exo JL, et al. Hemorrhagic shock shifts the serum cytokine profile from pro-to anti-inflammatory after experimental traumatic brain injury in mice. Journal of neurotrauma 2014; 31: 1386–1395. 2014/04/30. DOI: 10.1089/neu.2013.2985.

